# Regulation of trehalose biosynthesis and thermotolerance by the *Cryptococcus neoformans* HECT E3 ubiquitin ligase Rsp5

**DOI:** 10.64898/2026.01.07.698137

**Authors:** Alejandro L. Antonia, Lukas M. du Plooy, Siobhan R. Duffy, Jeffrey Kuhn, Erik J. Soderblom, J. Andrew Alspaugh

## Abstract

Microorganisms including fungi adapt to profound changes in their local environment during human infections. After exposure to high temperature and other stress conditions, the opportunistic fungal pathogen *Cryptococcus neoformans* enacts changes in metabolism, cell wall structure, and transmembrane transport that allow it to survive and proliferate in a mammalian host. This stress response program is regulated by the HECT E3-ubiquitin ligase Rsp5 which is required for growth at high salinity, pH and temperature. However, the complete set of Rsp5 substrates that direct these molecular changes remains incompletely understood. Here we demonstrate that *C. neoformans* Rsp5 confers increased tolerance to temperature and salt stress in part through regulation of the trehalose biosynthesis pathway. Two enzymes in the trehalose biosynthesis pathway, Tps1 and Tps2, are differentially ubiquitinated by Rsp5 after exposure to stress conditions. We directly measured trehalose production after exposure to high temperature and found that a *C. neoformans* strain lacking Rsp5 is unable to induce trehalose production. Quantitative proteomic analysis of the *C. neoformans* response to high salinity identified Rsp5-dependent and independent adaptations to osmotic stress, and that Rsp5-dependent ubiquitination does not alter the abundance of Tps1 or Tps2. These results demonstrate that regulation of trehalose biosynthesis is one of the cellular mechanisms by which Rsp5-dependent ubiquitination in *C. neoformans* facilitates survival in response to stressors encountered in the human infection environment.

**Importance:** *Cryptococcus neoformans* is an opportunistic fungal pathogen that kills over 180,000 people every year with few effective treatment options. As a yeast that normally lives in the environment, *C. neoformans* has to survive large changes in its physical environment, including elevated body temperature, when it causes human infections. Here we show how *C. neoformans* uses a protein modification to regulate production of a fungal-specific metabolic pathway important for survival at human body temperature. Unraveling how environmental fungi tolerate and survive temperature and other stressors will help to understand how they cause disease and identify new and better ways to treat these deadly infections.

## Introduction

Microorganisms often encounter drastic shifts in their physical environment requiring them to alter their metabolism and cellular structures. For the environmental and pathogenic yeast *Cryptococcus neoformans,* the ability to successfully navigate the transition from surviving in soil and trees to a mammalian host allows this microorganism to cause disease, resulting in over 100,000 deaths annually primarily in immunocompromised individuals(1–3). In response to stressors in human hosts such as elevated temperature and pH, *C. neoformans* undergoes profound molecular adaptations that allow it to survive and proliferate in this hostile environment(4–7).

Host-derived stress adaptations are coordinated by several central signaling pathways. These allow for survival at elevated temperatures, variations in pH, and in the presence cell wall stressors (8–11). For example, the *C. neoformans* high-osmolarity glycerol (Hog) response pathway consists of a two-component system in which membrane bound proteins sense environmental changes and activate an intracellular kinase signaling cascade which allows the yeast to tolerate cell wall, high temperature, and osmotic stressors(12, 13). Recent advances have also highlighted that post-translational modifications regulate *C. neoformans* stress adaptation. For example, in response to infection-related stress conditions, targeted protein ubiquitination can direct alterations in cell membrane structure through inositol sphingolipid biosynthesis and cell wall structure through chitin production(14, 15). Ubiquitin is a 76 amino acid moiety which is post-translationally covalently linked to specific lysine residues in target proteins. The ubiquitin cascade consists of E1 activating enzymes, E2 conjugating enzymes, and E3 ligases. The E3 ligases are the most numerous and diverse components of this cascade and thus help determine ubiquitin target substrate specificity.

The E3-ubiquitin ligase Rsp5 in *C. neoformans* is required for growth under host-relevant stress conditions. When the corresponding gene is deleted, this pathogen becomes avirulent in animal models of infection(15). We have previously demonstrated under high salinity and alkaline growth conditions that Rsp5 differentially ubiquitinates over 180 proteins. A detailed analysis of proteins with known Rsp5 binding motifs highlights the pleiotropic activity of this enzyme with its substrates involved in transmembrane transport, cell wall structure, and transcriptional responses to temperature stress(15). Here a comparative analysis between substrates ubiquitinated differentially between the high salinity and alkaline stress conditions identified a novel mechanism for Rsp5 in regulating trehalose biosynthesis.

Trehalose is a disaccharide with a biosynthetic pathway conserved across the fungal kingdom including pathogenic fungi, which is involved in maintenance of cell wall structure, tolerance to temperature stress, sexual differentiation, and as an alternative energy source(16, 17). For these reasons, disruption of the *C. neoformans* trehalose biosynthesis pathway leads to impaired virulence(18). The biosynthesis of trehalose in fungi including *C. neoformans* canonically includes two enzymes: 1) Tps1 which converts glucose-6-phosphate to trehalose-6-phosphate and 2) Tps2 which converts trehalose-6-phosphate to trehalose. Because trehalose biosynthesis is required for fungal virulence but is not present in mammals, it represents an enticing novel target for anti-fungal development(19). However, the precise mechanisms of regulation of the trehalose biosynthesis pathway by post-translational ubiquitination, especially in regard to microbial pathogens, has yet to be fully described.

In this study we explore the relationship between Rsp5-mediated ubiquitination and trehalose biosynthesis. An unbiased analysis of Rsp5 target proteins identified trehalose biosynthesis proteins as possible targets of ubiquitination mediating microbial response to host-derived stresses. By mass-spectrometry based direct protein quantification we show that Rsp5 does not alter the total protein abundance of the known components of the trehalose biosynthesis pathway. Finally, we demonstrate that deletion of Rsp5 impairs *C. neoformans* induction of trehalose production in response to temperature stress, suggesting that Rsp5-dependent temperature sensitivity is in part mediated by osmotic tolerance. Together these studies are the first demonstration of post-translational modifications regulating trehalose biosynthesis in pathogenic fungi.

## Results

### *C. neoformans* Rsp5 ubiquitinates effectors of microbial stress response pathways including trehalose biosynthesis

*C. neoformans* is an opportunistic fungal pathogen that needs to adapt to significant changes in its biological and abiotic environment when moving between its environmental niche and human host. This is in part achieved through changes in ubiquitination. We have shown that the HECT E3-ubiquitin ligase Rsp5 is required for stress adaptation and virulence(15). Prior screens to identify changes in ubiquitination of specific substrates under high salt conditions and alkaline stress conditions led to mechanistic studies confirming that Rsp5 regulates both nucleobase transport and Rim101 pathway activation as two distinct downstream mechanisms of adaptation to host-stress conditions(15, 20). However, it remains unclear whether Rsp5-mediated changes in ubiquitination are a common response to host-derived stress in general or specific to individual stressors.

A comparison of differentially ubiquitinated substrates between the conditions of alkaline stress and high salinity suggests that Rsp5 has a set of substrates that are generally altered under host-stress conditions and a set of substrates that are specific to unique individual stress conditions. The differential ubiquitination of substrates by Rsp5 was determined by incubating both wildtype *C. neoformans* and a *C. neoformans rsp5Δ* mutant strain under high salinity (NaCl 1.5M) or alkaline (pH 8.15) conditions for 1 hour as previously described(15, 20). We used a stringent threshold for enrichment of a 5-fold increase in ubiquitination between a wildtype strain and *rsp5*Δ strain to avoid interpreting false positive substrates. This identified 46 potential substrates under alkaline stress and 189 substrates under high salinity stress. Of these, 32 substrates were enriched as likely Rsp5 substrates under both stress conditions (Fig 1A).

**Figure 1.**
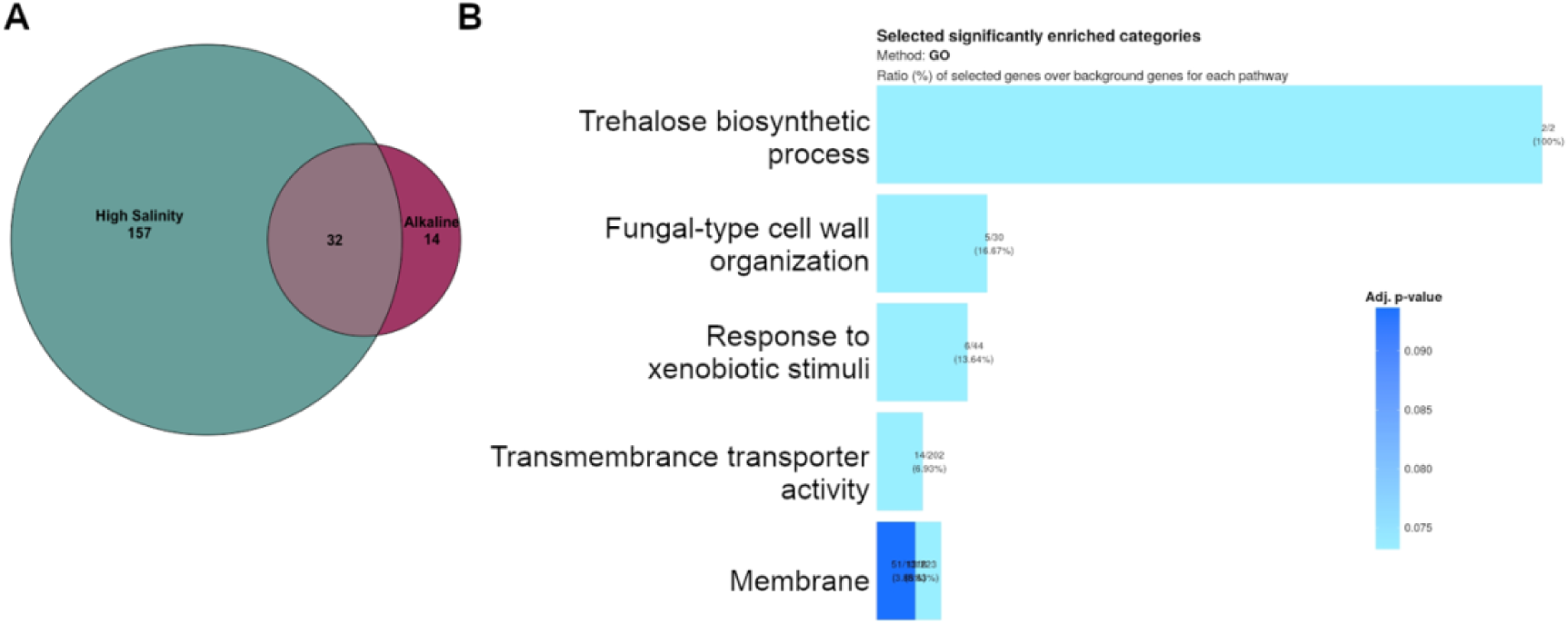
*C. neoformans* E3-ubiquitin ligase Rsp5 ubiquitinates distinct substrates in response to specific stress conditions, including the trehalose biosynthesis pathway. (A) Comparison of differential ubiquitin of Rsp5 substrates between host-derived stress conditions. *C. neoformans* was exposed to either high salinity (NaCl 1.5M) or alkaline pH (pH 8.15) for 1 hour prior to measuring ubiquitination of substrates by LC/MS-MS present in a wildtype but not in the *rsp5*Δ strain. B) Gene ontology analysis identified the trehalose biosynthesis pathway as significantly enriched among Rsp5 substrates after exposure to osmotic stress. The substrates exclusively ubiquitinated by Rsp5 under high salinity stress were analyzed for enrichment of assigned gene ontology terms using the FungiFun3 webserver.

To identify novel mechanisms of stressor-specific adaptations, we then performed gene ontology (GO) analysis for the set of proteins unique to either alkaline stress (no significant GO term enrichment observed) or high salinity conditions. The high salinity-specific Rsp5 substrates were enriched for those predicted to mediate phenotypes that we have previously described and characterized, including transmembrane transport and cell membrane integrity(15, 20). However, it also revealed that two enzymes in the trehalose biosynthesis pathway CNAG_05292 (Tps1) and CNAG_03765 (Tps2) are preferentially ubiquitinated by Rsp5 under high salt but not alkaline stress conditions (Fig 1B).

This suggested a novel link between Rsp5-dependent ubiquitination and trehalose biosynthesis, two central components of *C. neoformans* adaptation to host stress and virulence(15, 18–20). Thus we sought to investigate whether Rsp5-mediated ubiquitination changes the abundance of trehalose biosynthesis enzymes under host stress conditions, and whether this can provide insight into mechanisms of Rsp5-dependent thermotolerance.

### The *C. neoformans* proteomic response to high salinity includes Rsp5-dependent and independent changes

Because of the role of ubiquitination and thus Rsp5 in protein homeostasis during *C. neoformans* response to host-stress conditions, we performed whole cell proteomics to better characterize the molecular basis for the osmotic stress defect in this mutant strain. We subjected *C. neoformans* wildtype and *rsp5*Δ strains to high salinity (NaCl 1.5M) osmotic stress for 1 hour. Total protein lysates were harvested for quantitative mass spectrometry analysis to analyze changes in the proteome. Overall, 128 proteins were significantly changed using a permissive threshold of absolute fold change greater than 1.25 after exposure to high salinity stress (Fig 2A and Table S1). Of these 128 proteins, only 5 were previously identified as possible Rsp5 substrates (Fig 2A). Of these five genes, two have predicted roles in nucleobase transport (CNAG_00597 and CNAG_04632), two have predicted roles in cellular metabolism (CNAG_03040 and CNAG_04920), and one has no predicted function (CNAG_03415). Interestingly, of the two predicted amino acid transporters, CNAG_04632 (Nbt1) localization and abundance is altered in a Rsp5-dependent manner(15), and CNAG_00597 is transcriptionally upregulated in response to osmotic stress(21). Gene ontology analysis for the 128 proteins with changed abundance after osmotic stress did not identify any significantly enriched pathways. However, assigned functional terms for individual proteins with altered abundance in this assay highlight those with known roles in *C. neoformans* temperature and osmotic tolerance. For example, the protein Msb2 is increased in abundance by 7.4-fold (p=0.0003) after exposure to NaCl stress (Fig 2B). The homolog of this protein in *Saccharomyces cerevisiae* is a membrane sensor of osmotic stress that activates a kinase signaling cascade which promotes osmotic stress tolerance (22). In *C. neoformans,* Msb2 contributes to osmotic stress tolerance with other components of the HOG signaling pathway, and deletion leads to impaired survival of *C. neoformans* in a murine pulmonary infection model(13). Therefore, an analysis of the *C. neoformans* proteomic response to high salinity identifies both Rsp5-dependent and independent mechanisms of osmotic stress tolerance.

**Figure 2.**
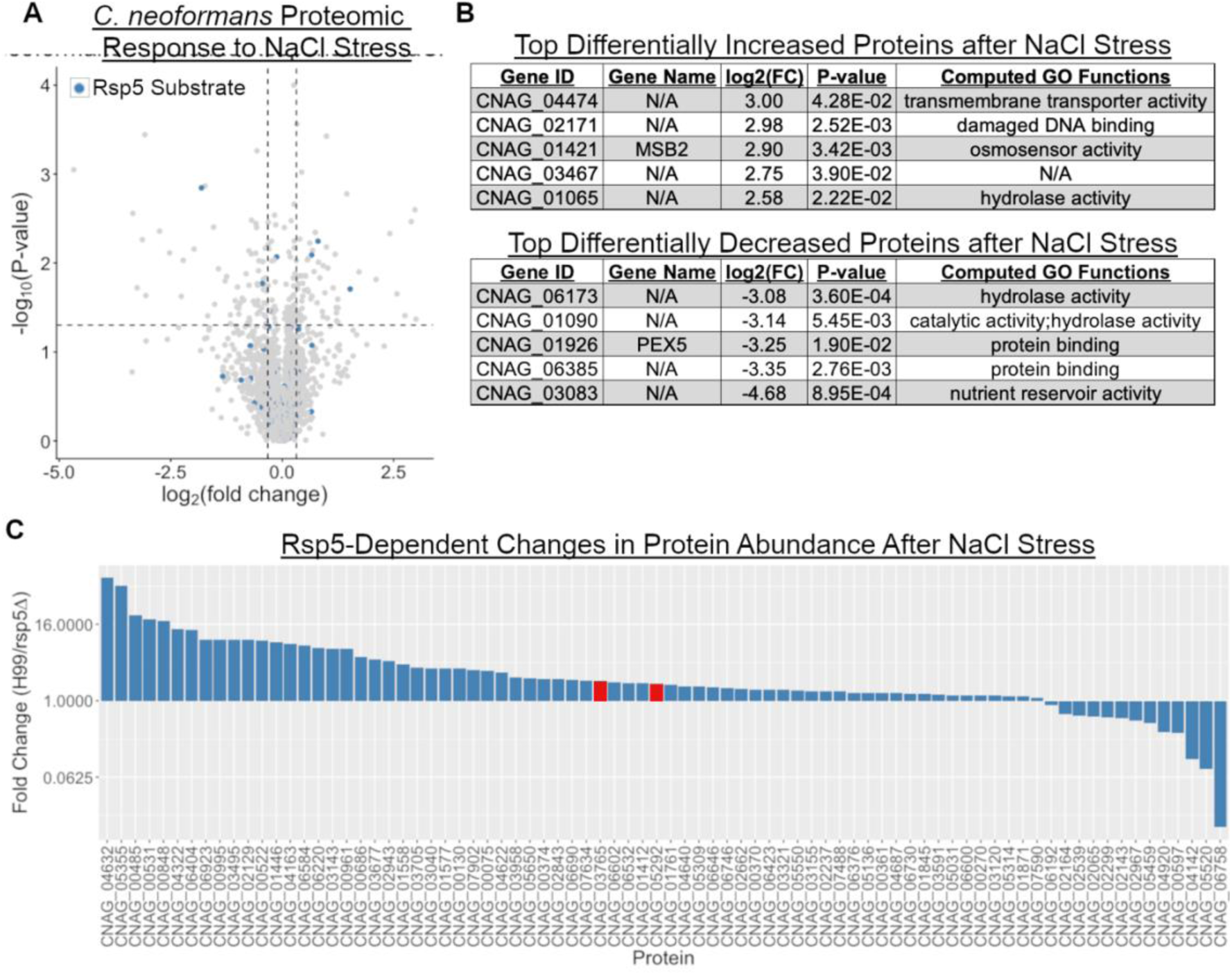
The *C. neoformans* proteome in response to osmotic stress includes both Rsp5-dependent and Rsp5-independent changes. (A) Volcano plot demonstrating *C. neoformans* changes in proteome in response to osmotic stress. Blue dots represent previously identified substrates of Rsp5 ubiquitination. Wildtype *C. neoformans* was incubated in liquid YPD supplemented with 1.5M NaCl for 1 hour prior to protein harvest. Protein quantification was performed by liquid chromatography-tandem mass spectrometry (LC-MS/MS). (B) The top 5 differential expressed proteins after osmotic stress include known components of the *C. neoformans* osmotic stress response such as Msb2. Proteins ranked by the fold change (FC) of wildtype *C. neorofmans* expression under NaCl stress compared to the control condition. (C) Rsp5-dependent changes in protein abundance after osmotic stress with 1.5M NaCl were determined by calculating the fold change between wildtype and *rsp5*Δ strains of *C. neoformans*, where lower fold-change represents proteins that have Rsp5 dependent decrease in protein abundance. Red bars represent known components of the trehalose biosynthesis pathway (Tps1 and Tps2).

### Rsp5 does not alter the total protein abundance of trehalose biosynthesis enzymes in response to osmotic stress

Post-translational ubiquitination can alter substrate total abundance, localization, or function. To test whether Rsp5 alters Tps1 or Tps2 abundance, we analyzed the subset of likely Rsp5 substrates (identified in Fig 1A) in the quantitative mass spectrometry dataset in response to high salinity. Rsp5-dependent changes in protein abundance were determined by comparing the fold change between the wildtype *C. neoformans* strain and the *rsp5*Δ mutant strain (Table S2). In total, 38 of the likely Rsp5 substrate proteins had at least a 2-fold decrease in abundance in the absence of Rsp5 which would be consistent with ubiquitin-mediated proteasomal degradation (Fig 2C). However, Tps1 (1.84-fold increase) and Tps2 (2.02-fold increase) did not have a decrease in total protein abundance in the absence of Rsp5. Therefore, it is likely that the Rsp5-mediated ubiquitination of trehalose biosynthesis enzymes involves proteasome-independent cellular processes in response to osmotic stress.

### Rsp5 ubiquitinates lysine residues distinct from the catalytic regions of Tps1 (lysine-228) and Tps2 (lysine-78)

Trehalose production is the result of an enzymatic cascade involving two enzymes: 1) Tps1 or alpha,alpha-trehalose-phosphate synthase (UDP-forming) and 2) Tps2 or Trehalose-6-phosphate phosphatase. Our prior differential ubiquitination assay for discovering Rsp5 substrates used liquid chromatography-tandem mass spectrometry (LC-MS/MS) to identify proteins present after stress conditions in either the *C. neoformans* wildtype or *rsp5Δ* strain after enriching with an antibody that recognizes the remnant ubiquitin stump after trypsin digestion(15, 20). One advantage of this approach is that the remnant of the anti-ubiquitin antibody can be resolved by LC-MS-MS, thus providing information on specific residues that Rsp5 likely ubiquitinates. We mapped these residues for both Tps1 (K-228) and Tps2 (K-78) and found that for both enzymes the ubiquitinated sites are distant from known catalytic or functional residues (Fig 2B)(19, 23). Interestingly, previous investigation has demonstrated that the location of the Tps1 K228 residue is located within an internal disordered domain (IDD) of uncertain functional significance, but which is exclusively found in basidiomycete fungi(19). Although the Rsp5 ubiquitinated residue on Tps2 (K78) is in a region without structural data, we observe a similar phylogenetic pattern for the complete Tps2 amino acid sequence as compared to the Tps1 sequences. Tps2 sequences from representative fungi distinctly cluster between the ascomycete and basidiomycete groups. Similar to Tps1, the basidiomycete Tps2 homologs contain several internal regions not present in ascomycetes. Also similar to Tps1, the ubiquitinated lysine (K78) on Tps2 is located in one such basidiomycete-specific region (Fig 2C). This suggests that Rsp5-mediated ubiquitination of these trehalose biosynthesis enzymes may be a unique phenomenon in this fungal phylum.

### *C. neoformans t*rehalose production after temperature stress is Rsp5-dependent

We then sought to directly measure how Rsp5 alters *C. neoformans* production of trehalose after temperature stress, the most well studied condition in which trehalose is cytoprotective. In the trehalose biosynthesis pathway, glucose-6-phosate is first converted to trehalose-6-phosphate, which is in turn converted to trehalose. If *C. neoformans* is unable to perform the first step in this enzymatic pathway, then the input glucose-6-phosphate is shunted to increase glycogen production (summarized in Fig 2A). We measured trehalose production after growth at 37°C for 8 hours in liquid media. At 37°C the *C. neoformans* wildtype strain produced over twice as much trehalose per cell compared to the *rsp5*Δ strain (2.4-fold increase; p=0.0002) (Fig 2D). We confirmed that this is an Rsp5-dependent effect by demonstrating complete restoration of trehalose production by complementation of *RSP5* into the mutant background. Overall, the wildtype and *rsp5*Δ+*RSP5 C. neoformans* strains significantly upregulate trehalose production after temperature stress while *rsp5*Δ, *tps1*Δ, and *tps2*Δ strains do not similarly induce trehalose (Fig 2D). Finally, we tested whether the lack of Tps1 activity or trehalose production results in glycogen accumulation by performing colorimetric iodine staining, a marker of total carbohydrate content, which has previously been validated as a surrogate for glycogen accumulation(24). Interestingly, the *rsp5*Δ strain did not accumulate glycogen after temperature stress (Fig 2E), suggesting that the Tps1 enzyme retains sufficient activity in the absence of Rsp5 to prevent hyperaccumulation of glycogen despite the notable decrease in overall trehalose production.

### *C. neoformans* Rsp5-dependent temperature sensitivity is partially due to impaired osmotic tolerance

*C. neoformans rsp5*Δ, *tps1*Δ, and *tps2*Δ mutant strains are all thermosensitive at human physiologic temperature (37°C) and all correspondingly have known reduced virulence in animal models(15, 18). However, only *rsp5*Δ and *tps1*Δ mutants have clear growth defects under osmotic and alkaline stress (Fig 3A). The overlapping phenotypes with effectors that have established roles in *C. neoformans* stress tolerance may also provide insight into the molecular mechanisms of Rsp5-dependent stress adaptation. For example, the phenotype of impaired growth under high temperature for trehalose biosynthesis mutants is in part due to impaired osmotic tolerance. Accordingly, the addition of an osmotic stabilizer such as sorbitol to the growth medium restores thermotolerance to the *C. neoformans tps1Δ and tps2*Δ strains(18). Similarly, we also observed that for the *C. neoformans rsp5*Δ mutant, the NaCl growth defect occurs in a dose-dependent manner consistent with NaCl acting as an osmotic stressor (Figure 3B). Therefore, we hypothesized that supplementation with sorbitol would also be able to restore growth of the *C. neoformans rsp5*Δ strain at elevated temperatures. Indeed, we found that adding exogenous sorbitol was sufficient to restore growth of the *rsp5*Δ strain at the elevated human physiologic of 37°C (Figure 3C). This discovery of an additional mechanism of Rsp5-dependent tolerance to temperature stress likely reflects the pleiotropic and multi-targeted nature of Rsp5 (Fig 1A).

**Figure 3.**
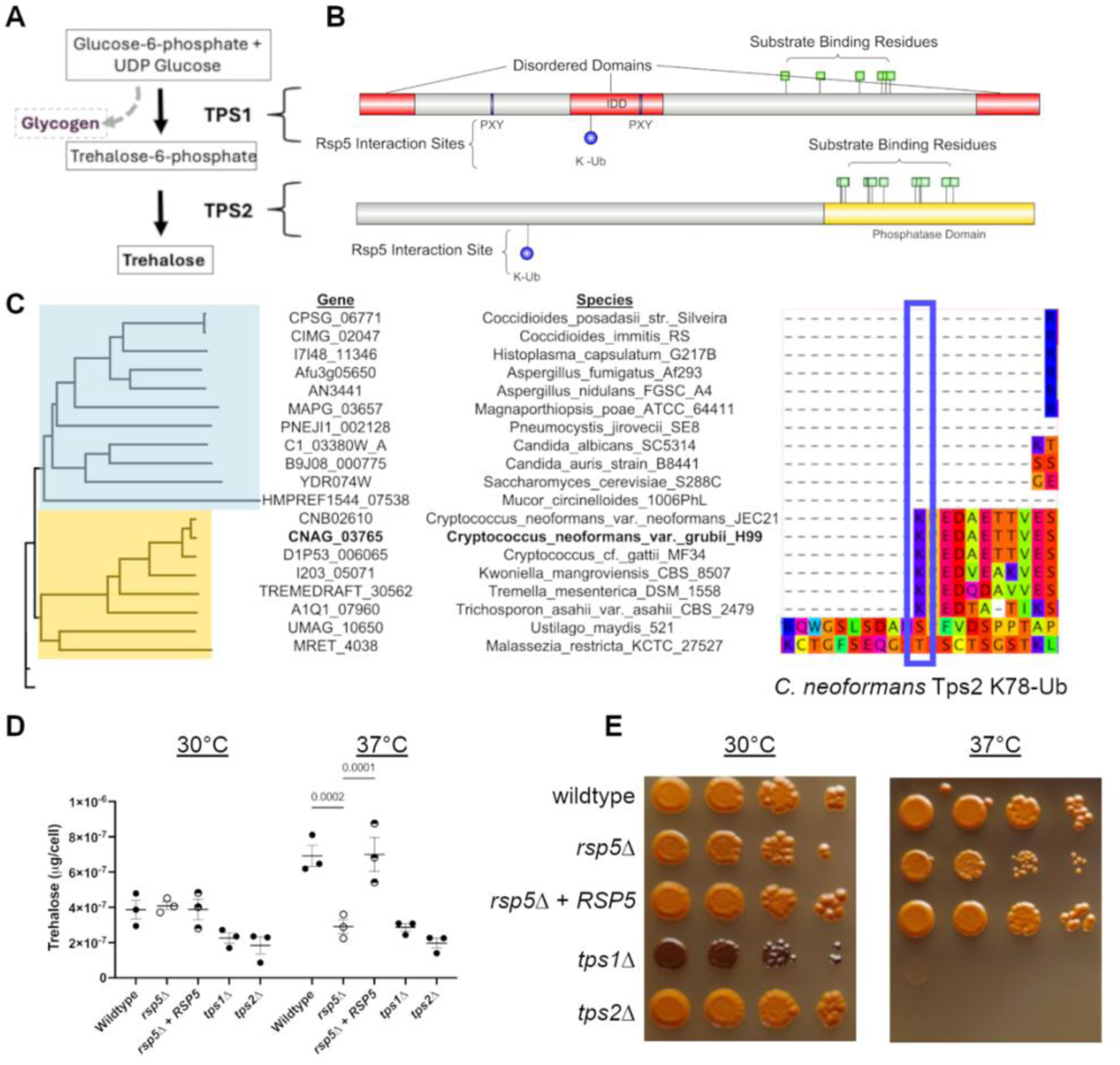
*C. neoformans* induction of trehalose production in response to temperature stress is Rsp5-dependent. (A) Simplified schema of trehalose biosynthesis including the enzymes alpha, alpha-trehalose-phosphate synthase (Tps1) and trehalose-6-phosphate phosphatase (Tps2). (B) Domain diagram of *Cn* Tps1 and Tps2. Rsp5 interaction sites are highlighted in blue including putative PXY binding motifs and the ubiquitination sites K228 on Tps1 and K78 on Tps2 identified by LC-MS-MS after exposure to high salinity stress. Known enzymatic residues are highlighted by green squares. (C) The Tps2 K78 residue ubiquitinated by Rsp5 is in a basidiomycete specific segment of the protein. Tps2 homologues were identified by BlastP with search limited to a manually curated set of representative reference organisms. The phylogeny and multisequence alignment were generated by Clustal algorithm. Clades that included proteins with known alternative functions (such as Tps1) were collapsed in the presented phylogeny. (D) After temperature stress *Cn* upregulation of trehalose production is Rsp5-dependent. Trehalose quantification determined after 8 hours at indicated temperature and normalized per individual cell as quantified by hemocytometer. Mean and standard error of the mean plotted from 3 biological replicates. P-values calculated by two-way ANOVA with Tukey’s post-hoc test. (E) Loss of Rsp5 does not result in hyperaccumulation of glycogen. Each strain was diluted to a starting OD of 1.0 with 4 serial 10x dilutions prior to pinning onto YPD agar plates and incubating at the indicated temperature for 72 hours. Iodine vapor was used to stain each plate for glycogen with violet color representing greater concentration of carbohydrates including glycogen. Representative images shown from three separate biological replicates.

## Discussion

*C. neoformans* utilizes the E3 ubiquitin ligase Rsp5 to adapt to a range of host-derived stressors including high salinity, pH, and elevated temperature. By comparing the differences in ubiquitinated protein substrates between high salinity and alkaline stress conditions, we describe that Rsp5 ubiquitinates a core set of substrates in response to general cell stress in addition to ubiquitination of stress condition-specific substrates. The high salinity stress-specific Rsp5 substrates include trehalose biosynthesis enzymes, a *C. neoformans* stress response pathway with known roles in temperature sensitivity. Thus, we describe for the first time in a pathogenic fungus that ubiquitination regulates the trehalose biosynthesis pathway in response to host-derived stress conditions and identify an additional mechanism underlying Rsp5-dependent temperature tolerance.

However, the mechanism by which Rsp5 integrates distinct environmental cues to coordinate context-specific downstream adaptations remains unknown. There are several potential alternative models that should be investigated. First, under specific stress conditions the subcellular localization of Rsp5 may be altered and thereby place it in the proximity of relevant substrates that are then differentially ubiquitinated. Second, Rsp5 may retain the same substrate affinity in different stress conditions but the relative abundance of its substrates is increased either transcriptionally or translationally during stress, thereby resulting in differential ubiquitination. Either of these alternative hypotheses could be further influenced by changes in Rsp5-adaptor proteins that coordinate the interaction between Rsp5 and many of its substrates. Further understanding of how Rsp5 responds to distinct stressors is likely to yield mechanistic insight into critical *C. neoformans* stress response pathways.

One of the most important ways in which ubiquitin ligases control substrate specificity is the use of differentially targeted adaptor proteins. For example, the arrestin-like Ali1 protein directs enzymes involved in fatty acid synthesis to sites of rapid cell membrane expansion during budding(25). A distinct arrestin domain-containing protein, Ali2, mediates a separate subset of

Rsp5 interactions with target proteins involved in alkaline stress response(15). Despite demonstrating that *C. neoformans* Rsp5 is required for optimal trehalose biosynthesis in response to high temperature stress, future studies will be necessary to elucidate how Rsp5 interacts with components of the trehalose biosynthesis pathway and whether additional adaptor proteins support those interactions. It is possible that Tps1 and Tps2 directly bind to Rsp5 prior to their ubiquitination. These E3 ubiquitin ligase-target protein interactions are often facilitated by internal proteins sequences such as the PXY motif required for Rsp5 interactions with its targets. We note that Tps1 has one canonical Rsp5 binding PXY motif. This suggests that Rsp5-Tps1 interactions may occur independently from adaptor proteins. In contrast, Tps2 does not contain any such motifs. Interestingly, our group has previously identified Tps2 as physically interacting with the Rsp5 adaptor protein Ali1(15, 25) suggesting a model for the Rsp5-Tps2 interaction through this adaptor protein. Downstream of the ubiquitination event, the direct impact of Rsp5-dependent ubiquitination of Tps1 and Tps2 needs to be further elucidated. We demonstrated by direct measurement of Tps1 and Tps2 that Rsp5 dependent ubiquitination does not alter the total abundance of these substrates at either permissive or stress conditions (Fig 4C.) In other model organisms Rsp5-dependent ubiquitination events frequently result in changes in subcellular localization by differential vesicular sorting(26) suggesting a likely alternative model for how *C. neoformans* Rsp5 contributes to trehalose production in response to host stressors.

**Figure 4.**
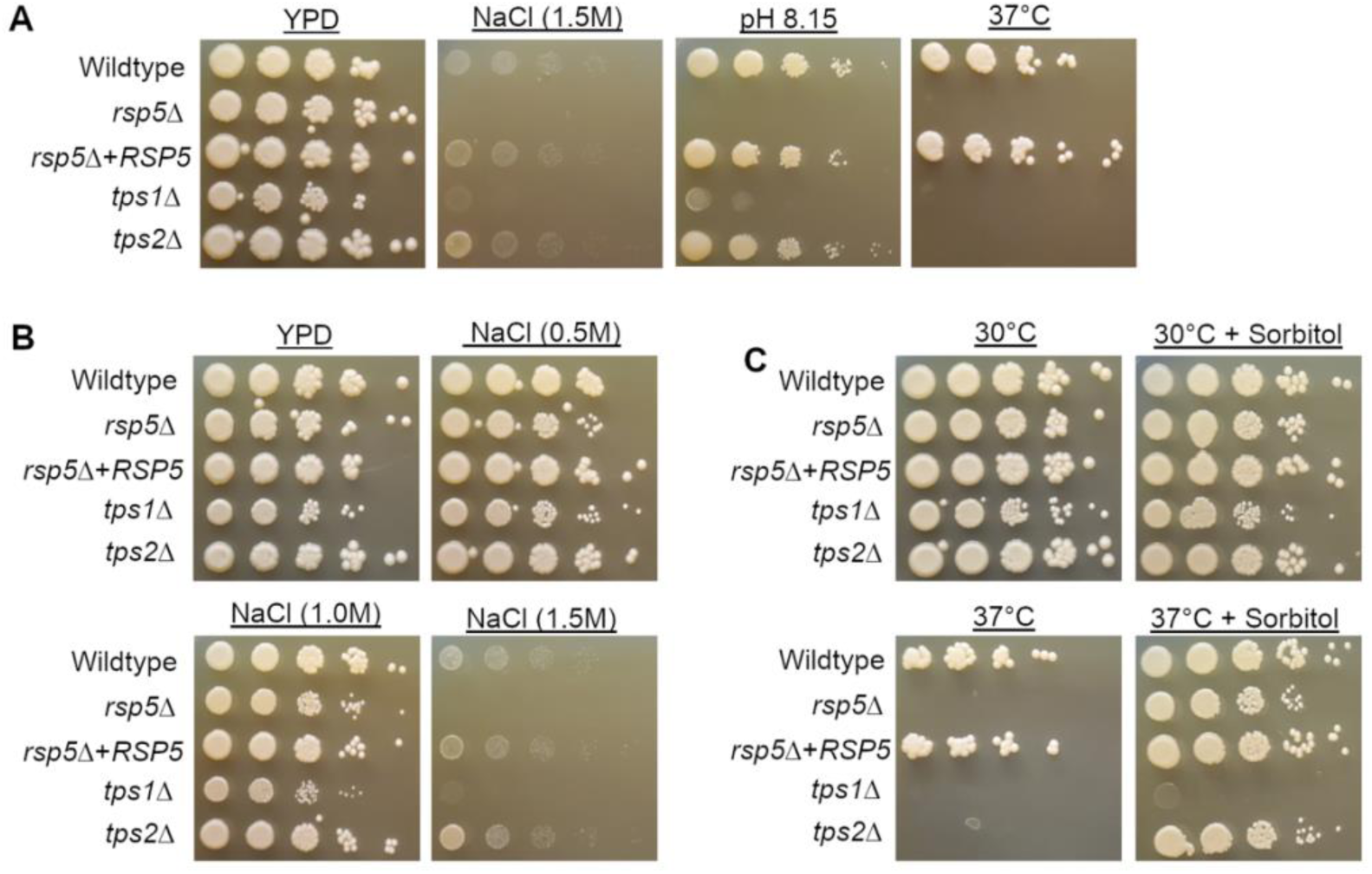
Rsp5-dependent temperature sensitivity is partially due to osmotic stress tolerance. (A) *C. neoformans rsp5*Δ and *tps1*Δ mutants have impaired growth under high salinity (NaCl 1.5M), alkaline (pH 8.15), and high temperature (37°C) stress conditions ; whereas the *tps2*Δ mutant only has a temperature-sensitive growth defect. (B) The temperature sensitivity of Rsp5 increases with dose-dependent increase in NaCl consistent with a model of osmotic stress. (C) The temperature-sensitive growth defect at 37°C for the *C. neoformans rsp5*Δ strain is partially restored by addition of the osmotic stabilizer sorbitol (1M). For (A-C), spot dilution assays performed by normalizing all strains to OD600 of 1.0 followed by serial 10x dilutions prior to pinning onto YPD plates with the indicated additive. Plates were then incubated at either 30°C or 37°C for 72 hours. Representative images shown from three separate biological replicates.

Additionally, for both enzymes in the trehalose biosynthesis pathway that are ubiquitinated by Rsp5, we identified the ubiquitination event to occur on a lysine residue located in a basidiomycete-specific region of the protein sequence. *C. neoformans* is in the basidiomycete fungal phylum, which also contains the major plant pathogen *Ustilago maydis*, but which is divergent from ascomycete fungi such as *Saccharomyces cerevisiae.* This suggests a hypothesis that in basidiomycete fungi, such as *C. neoformans,* trehalose biosynthesis enzymes have undergone selective pressure to fine-tune the post-translational regulation of the enzymes rather than selection to alter enzyme function or specificity. Future studies will be needed to test whether the relationship between Rsp5 and trehalose biosynthesis has undergone convergent evolution in other fungi, and whether similar patterns have occurred in other Rsp5 substrates.

Here we demonstrate that the pleiotropic nature of the E3-ligase Rsp5 enzyme can facilitate *C. neoformans* stress adaptation coordinated to specific environmental cues. One of these stressor-specific mechanisms is through upregulation of trehalose biosynthesis. An improved understanding of the cross-talk between these key regulators of *C. neoformans* stress response will help us to better understand how this environmental yeast becomes an opportunistic pathogen.

## Materials and Methods

### Strains, media, and growth conditions

All strains used in this study were created in the *Cryptococcus neoformans* var *grubii* H99 background (Table 1) (27). The *C. neoformans tps1*Δ and *tps2*Δ strains were provided as gifts from the lab of John R. Perfect and validated by presence-absence polymerase chain reaction (PCR) with the following primer pairs: Tps1_Fwd (ATGCACATGGGCGGAACATA) with Tps1_Rev (GCCAGTTCCGACTAAAGCCT) and Tps2_Fwd (GCGGATCAGAAAGAGATTGC) with Tps2_Rev (CTTCTGATAGGCCAGCCAAG). Strains were recovered from glycerol stocks stored at -80°C onto solid YPD media (1% yeast extract, 2% peptone, 2% dextrose, and 2% agar) at 30°C. For each experiment strains were grown in liquid YPD media at 30°C in a shaking incubator at 150rpms unless otherwise stated. Liquid YPD media was maintained at pH 5.5 to 6.0 unless otherwise indicated. For alkaline stress media, YPD media was buffered with 150mM HEPES and then adjusted to pH 8.15 with potassium hydroxide. This pH that is slightly more alkaline than host pH (7.4) was empirically chosen to best visualize and distinguish cellular phenotypic differences between acidic and alkaline pH(5, 20). For high salinity stress media, the indicated concentration of sodium chloride was added to each media condition.

**Table 1:** *Cryptococcus neoformans* strains used in this study.

| Strain name/number | Genotype | Reference/Source |
| --- | --- | --- |
| H99 | *MATα* | (Perfect et al., 1980) |
| MDP01 | *MATα rsp5Δ::NAT* | (Du Plooy et al. 2024) |
| MDP26 | *MATα rsp5Δ::NAT+RSP5* | (Du Plooy et al. 2024) |
| Tps1Δ | *MATα tps1Δ::NAT* | (Washington et al. 2024) |
| Tps2Δ | *MATα tps2Δ::NAT* | Gift from John R. Perfect |

### Proteomics and differential ubiquitination determination

To assay changes in total protein abundance after the osmotic stress of high salinity, after 18 hour overnight culture the indicated *C. neoformans* strains were diluted to an OD_600_ of 3.0 in 4mL of YPD buffered to pH 4.0 with or without the addition of 1.5M sodium chloride (NaCl). Strains were incubated at 30°C for 1 hour prior to protein harvest. Protein harvest was performed by washing the cell pellet with distilled water twice, then resuspending in urea lysis buffer (8M urea and 50mM NH_2_HCO_3_) supplemented with cOmplete mini, EDTA-free protease inhibitor (Roche) Pierce phosphatase-inhibitor mini tablets (Thermo Scientific) and 1mM phenylmethylsulfonyl fluoride (PMSF). Cells were then lysed by bead beating and with total protein normalized prior to protein quantification by liquid chromatography tandem mass-spectrometry (LC-MS/MS) as previously described(15, 25).

LC-MS/MS analysis was performed with the assistance of the Duke Proteomics and Metabolomics Core Facility as previously described(15, 25). In brief, following protein solubilization in 8M urea/50 mM ammonium bicarbonate, samples were brought to 5% SDS and reduced with 10 mM dithiolthreitol for 15 min at 32C, alkylated with 25 mM iodoacetamide for 20 min at room temperature, then trypsin digested with sequencing grade trypsin (Promega) on an STrap Micro (Protify) cartridge using manufacturer recommended protocols.

Quantitative LC/MS/MS was performed on 250ng of sample using an EvoSep One UPLC coupled to a Thermo Orbitrap Astral high resolution accurate mass tandem mass spectrometer (Thermo). Briefly, each sample loaded EvoTip was eluted onto a 1.5 µm EvoSep 150μm ID x 8cm performance (EvoSep) column using the SPD60 gradient at 55°C. Data collection on the Orbitrap Astral mass spectrometer was performed in a data-independent acquisition (DIA) mode of acquisition with a r=240,000 (@ m/z 200) full MS scan from m/z 380-980 in the OT with a target AGC value of 4e5 ions. Fixed DIA windows of 4 m/z from m/z 380 to 980 DIA MS/MS scans were acquired in the Astral with a target AGC value of 5e4 and max fill time of 6 ms. HCD collision energy setting of 27% was used for all MS2 scans.

Raw data were imported into Spectronaut v20 (Biognosis) and MS/MS data was searched against a SwissProt *C. neoformans H99* database (downloaded 2024). A library free Direct DIA+ approach within Spectronaut was used to perform the database searches set at a maximum 1% peptide false discovery rate based on q-value calculations. Database search parameters included fixed modification on Cys (carbamidomethyl) with variable modification on Met (oxidation). Full trypsin enzyme rules were used along with 10ppm mass tolerances on precursor ions and 20ppm on product ion. Peptide homology was addressed using razor rules in which a peptide matched to multiple different proteins was exclusively assigned to the protein with higher % sequence coverage. A MaxLFQ rollup strategy was deployed to roll up from the precursor level to the protein level(28). Determination of differentially ubiquitinated substrates was described previously for high salinity (NaCl 1.5M) and alkaline (pH 8.15) stress(15, 20). In brief, 1mg of trypsin digested cell lysate was enriched using the PTMScan® HS Ubiquitin/SUMO Remnant Motif (K-ε-GG) Kit (Cell Signaling Technologies) according to manufacturer protocols (see https://media.cellsignal.com/pdf/59322.pdf). Eluted peptides were analyzed using a nanoAcquity UPLC system (Waters Corp) coupled to a Thermo Orbitrap Fusion Lumos high resolution accurate mass tandem mass spectrometer (Thermo). Peptides were trapped on a Symmetry C18 20 mm × 180 µm trapping column (5 μl/min at 99.9/0.1 v/v water/acetonitrile), after which the analytical separation was performed using a 1.8 µm Acquity HSS T3 C18 75 µm × 250 mm column (Waters Corp.) with a 90-min linear gradient of 5 to 30% acetonitrile with 0.1% formic acid at a flow rate of 400 nanoliters/minute (nL/min) with a column temperature of 55C. Data collection was performed in a data-dependent acquisition (DDA) mode of acquisition with a r=120,000 (@ m/z 200) full MS scan from m/z 375 – 1500 with a target AGC value of 4e5 ions was performed. MS/MS scans were acquired in the linear ion trap in “rapid” mode with a target AGC value of 1e5 and max fill time of 100 ms.

Data were imported into Proteome Discoverer 3.0 (Thermo Scientific Inc.) and the MS/MS data was searched against the SwissProt *C. neoformans H99* database (downloaded in 2023) and an equal number of reversed-sequence “decoys” for false discovery rate determination. Sequest (v 3.0, Thermo PD) was utilized to produce fragment ion spectra and to perform the database searches. Database search parameters included fixed modification on Cys (carbamidomethyl) and variable modification on Met (oxidation) and Lys (-GG). Search tolerances were 2ppm precursor and 0.8Da product ion with full trypsin enzyme rules. Peptide Validator and Protein FDR Validator nodes in Proteome Discoverer were used to annotate the data at a maximum 1% protein false discovery rate based on q-value calculations. In this study, the set of substrates with at least 5-fold enrichment in the wildtype compared to *rsp5*Δ strain were used for further analysis. All mass spectrometry data has been uploaded to ProteomeXchange with the identifier PXD071777.

### Gene ontology analysis

Gene ontology analysis was performed using the FungiFun3 webserver (29). Overrepresentation analysis using gene ontology terms with *C neoformans* var grubii serotype A (strain H99/ATCC208821/CBS10515/FGSC9487) specified as reference species. Unless otherwise specified default parameters were used.

## Phylogenetic analysis

*C. neoformans TPS2 (CNAG_03765*) homologs were identified by BlastP search with E-value cutoff of 0.0001. The search was restricted to the following set of manually curated representative fungi: *Aspergillus fumigatus Af293, Aspergillus nidulans FGSC A4, Candida albicans SC5314, Candida auris strain B8441, Coccidioides immitis RS, Coccidioides posadasii str. Silveira, Cryptococcus cf. gattii MF34, Cryptococcus gattii VGII R265, Cryptococcus neoformans var. grubii H99, Cryptococcus neoformans var. neoformans JEC21, Histoplasma capsulatum G217B, Kwoniella mangroviensis CBS 8507, Magnaporthiopsis poae ATCC 64411, Malassezia restricta KCTC 27527, Mucor circinelloides 1006PhL, Pneumocystis jirovecii SE8, Saccharomyces cerevisiae S288C, Tremella mesenterica DSM 1558, Trichosporon asahii var. asahii CBS 2479, Ustilago maydis 521.* This resulted in identification of 64 proteins, which included the related but distinct protein Tps1. To restrict the phylogeny to Tps2 homologues the initial set of results was filtered to include only those with >40% sequence identity to the CNAG_03765 reference sequence. Multisequence alignment was then performed with default parameters using the Clustal algorithm on the EMBL-EBI webserver(30). The corresponding phylogenetic tree alignment was visualized in R using the ggtree package(31). Multisequence alignment was visualized with Jalview(32).

### Stress response assays

*C. neoformans* stress response phenotypes were assayed by spot-dilution assay as described previously(33). In brief, overnight cultures of the indicated strains of *C. neoformans* were diluted to an OD_600_ of 1.0 prior to making 5 serial 10x dilutions in 200μl of YPD in 96-well plate format. They were then spotted in replicate onto the indicated plate with a standardized volume using metal-pronged pinner. Plates were incubated in temperature controlled incubator at 30°C unless otherwise indicated.

### Trehalose quantification and glycogen staining

To measure intracellular trehalose concentration, after the indicated stress conditions cell concentration was determined using a hemocytometer. A standardized volume for the same total number of cells was centrifuged at room temperature at 3,000g for 5 minutes, and washed once with sterile deionized H_2_O. Then, the supernatant was removed and the remaining cell pellet was placed in a lyophilizer overnight. Cells were then disrupted by placing 3mm glass beads into the 15mL culture tube with the dried cell pellet followed by vortexing vigorously for 30 seconds. Cellular debris was then resuspended in 200μL of sterile PBS prior to transfer to a microcentrifuge tube. The cellular debris was then centrifuged in table top centrifuge at 13,000rpm for 2 minutes, and the supernatant transferred to new sterile microcentrifuge tube. Each sample was then treated overnight at 37°C with trehalase by combining 30μL of sample, 30μL of sodium citrate 0.1M, and 10μL of porcine trehalase at 0.00033U/mL working concentration (Millipore Sigma; T8778). A vehicle control was prepared in parallel for each sample with no added trehalase. Finally, the total glucose resulting from each sample was quantified using a colorimetric glucose detection assay per manufacturers protocol in microplate format (Millipore Sigma; GAGO20). Total trehalose was calculated as follows: 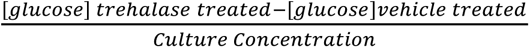. Culture concentration was determined by hemocytometer.

## Data availability

Data from the proteomics experiment is available in supplemental material and in ProteomeXChange database. All other data is available upon request.

## Acknowledgements

This work was supported by NIH RO1 grant AI175711 and R21 grant AI153799 to JAA. ALA was supported by the NIH funded Tri-Institutional Molecular Mycology and Pathogenesis Training Program (5T32AI052080). We thank Erica Washington, Jennifer Tenor, and John Perfect for thoughtful discussion of the assessment and sharing reagents related to *C. neoformans* trehalose-dependent stress response.

